# Cross-Modal Cue Effects in Motion Transparency Processing

**DOI:** 10.1101/242214

**Authors:** G.M. Hanada, J. Ahveninen, F.J. Calabro, A. Yengo-Kahn, L.M. Vaina

**Affiliations:** Brain and Vision Research Laboratory, Department of Biomedical Engineering, Boston University, Boston, MA, USA; Harvard Medical School – Athinoula A. Martinos Center for Biomedical Imaging, Department of Radiology, Massachusetts General Hospital, Charlestown, MA, USA; Department of Psychiatry and Department of Bioengineering, University of Pittsburgh, Pittsburgh, PA, USA

**Author notes:** Corresponding Author:* Jyrki Ahveninen, Ph.D. MGH/MIT/HMS-Martinos Center Bldg. 149 13^th^ Street, Charlestown MA 02129.

## Abstract

The everyday environment brings about many competing inputs from different modalities to our sensory systems. The ability to filter these multisensory inputs in order to identify and efficiently utilize useful spatial cues is necessary to detect and process the relevant information. In the present study, we investigate how feature-based attention affects the detection of motion across sensory modalities. We were interested to determine how subjects use intramodal, crossmodal auditory, and combined audiovisual motion cues to attend to specific visual motion signals. The results show that in most cases, both visual and auditory cues enhance feature-based orienting to a visual motion pattern that is presented among distractor patterns. Furthermore, in many cases, detection of transparent motion patterns was significantly more accurate after combined visual-auditory than unimodal attention cues. Whereas previous studies have shown crossmodal effects of spatial attention, our results demonstrate a spread of crossmodal feature-based attention cues, which have been matched for the detection threshold of the visual target. These effects were evident in comparisons between cued and uncued conditions, as well as in analyses comparing the effects of valid vs. invalid cues.

## Introduction

In everyday life, we are surrounded by scenes cluttered with many different static and dynamic objects, which cannot be processed simultaneously. Functioning in such environment requires attentional mechanisms whose effectiveness is increased by the availability of relevant unimodal and crossmodal sensory cues. In many cases, the unimodal cues are not from the principal modality of the target stimulus, such as when a looming sound guides visual attention to a particular object in a scene that is approaching instead of receding. Much of the previous research on motion detection and orienting of visual attention to motion patterns has, however, concentrated on unimodal studies.

Multisensory topics such as the effects of crossmodal spatial (Spence et al., 1998, Jack and Thurlow, 1973, Kopco et al., 2009) or temporal (Shams et al., 2000, Shams et al., 2002, Vroomen and de Gelder, 2000) cues on stimulus detection have been studied extensively. The spatial domain of multisensory perception is generally believed to be dominated by visual information (Jack and Thurlow, 1973, Kopco et al., 2009). However, auditory stimuli may provide coarser spatial orientation cues that guide the observers’ fine-grained visual attention to the relevant location in a stimulus-driven fashion and subsequently improve the perceptual performance (Beer and Röder, 2005, Ward et al., 2000, Driver, 2004, Driver and Spence, 1998).

In addition to the relatively large number of studies regarding spatial or temporal crossmodal influences, a small number of studies have examined crossmodal modulation of the processing of motion direction cues. These studies have suggested that selective attention to auditory or visual motion direction also spreads to the other modality (Beer and Röder, 2004), analogous to the effects of stimulus-driven spatial attention (Driver and Spence, 1998). There is also evidence that crossmodal cues support both auditory and visual motion perception (Cappe et al., 2009, Soto-Faraco et al., 2003, Schmiedchen et al., 2012), and modulate brain activations during motion discrimination tasks (Lewis and Noppeney, 2010, Kayser et al., 2017). Our recent studies also demonstrate that congruent auditory cue can help identify a moving object in an environment where the observer is in self-motion (Calabro and Vaina, 2011, Roudaia et al., 2018), a process highly important for our daily activities.

However, overall, the evidence regarding auditory influences on visual motion processing is not entirely consistent. Some studies suggest that while the crossmodal effects are observable in both directions, visual motion cues have larger influences on auditory motion perception than vice versa (Bertelson and Radeau, 1981). In the case of apparent motion illusion, one study reported that auditory cues produce no crossmodal effects while visual cues modulate the auditory apparent illusion very clearly (Soto-Faraco et al., 2004). There are also studies suggesting that combined multisensory information direction produces only small additional benefits in the discrimination of linear translational motion (Alais and Burr, 2004). It has been also argued that the effects of crossmodal auditory motion cues bias the post-perceptual decision making instead of modulating the sensitivity of visual detection of random dot motion direction, per se (Meyer and Wuerger, 2001).

In the present study, our goal was to determine how feature-based attention affects the detection of motion across sensory modalities. We investigated how subjects use visual, auditory, and audiovisual motion cues to attend to specific visual motion signals. The results show that in most cases, both visual and auditory cues enhance feature-based orienting to a visual motion pattern that is presented among distractor patterns.

## Material and Methods

### Participants

Twelve observers participated in the study (mean age = 24.75 years, SD = 4; all male). All had normal hearing and normal or corrected-to-normal vision. Three of the observers were authors; the rest of them were naïve as to the purpose of the experiment. All observers gave informed consent according to the Boston University Institution Review board. Prior to enrollment subjects underwent a rigorous training and practice for discrimination accuracy. Subjects were required to score more than 60% correct across all practice blocks otherwise they were excluded from the study. All subjects were able to achieve the required level of accuracy.

### Display and procedure

Participants were seated at 60 cm viewing distance from the computer monitor in a dark room and were adapted for 5 min to the background luminance of the monitor with head position stabilized with a chin and forehead rest. All stimuli were generated on a Mac Pro running MATLAB using the BRAVI-shell software developed in our laboratory based on the Psychophysical Toolbox (Brainard 1997 and Pelli 1997), and were presented on a 23” Apple LCD Cinema Display. All auditory cues were presented with Sennheiser HD201 headphones. We used a Minolta LS-100 light meter for monitor luminance calibration and a Scantek Castle GA-824 Smart Sensor SLM for acoustic calibration.

The visual stimulus consisted of Random Dot Kinematograms (RDK), 50 white dots, 43.9 cd m^−2^ luminance, shown on a gray background (9.9 cd m^−2^, and dot to background contrast was 23%), and moving at 5°/second. RDKs were presented in four circular apertures (8° diameter), each displayed within one of four quadrants of the computer screen, and with the center of each aperture being 8° from the fixation mark. Each aperture displayed transparent motion defined by two superimposed RDKs: in one the dots moved horizontally (0° or 180°) and in the other, they moved vertically (90° or 270°). Each RDK was set at 80% coherence such that for any given pair of frames, 80% of the dots displayed were moving in the selected direction (signal dots) while 20% of the dots were repositioned at random locations within the aperture (noise dots). Coherently moving dots were wrapped around the edge of the aperture to maintain a constant density throughout the stimulus duration.

In every trial, the horizontal direction within each aperture was randomly selected (e.g. 0°) to be the “target” direction and one of the four apertures was randomly selected to display the RDK with its horizontally moving dots in the “target” direction. The direction of motion in the other three apertures was opposite to the direction of the “target” (e.g. 180°). All four apertures simultaneously contained a superimposed second RDK of the same density and luminance as the first, but the motion was vertical (90° or 270°). In each aperture, the specific vertical direction was randomly selected (90° or 270°) and thus did not depend on the horizontally moving RDK in that aperture. Subjects were instructed to selectively attend to the specified plane of motion (e.g., horizontal). They were first asked to identify the target aperture (by pressing a predesignated button on the computer keyboard that indicates the quadrant where the target aperture is displayed). Immediately after this, they were asked to report the direction of the vertical motion in the target aperture (by pressing a predesignated button on the computer keyboard).

### Cues

Four types of cues were used to assess the effect of within- and cross-modal attention on facilitating the ability to identify the target aperture. All cue types provided horizontal motion direction information to the subjects prior to the start of the stimulus in which the horizontal motion direction would match the target horizontal motion direction during the stimulus display. In 20% of the trials (“invalid cue” trials), the horizontal motion direction of the cue was opposite to the direction in the target aperture.

1. No Cue: The aperture presented at the beginning of each trial remained blank for its duration and observers performed the task as described but without any cue. Subjects had to identify the target aperture based on the oddball motion direction (e.g., the single aperture with leftward motion, among three others moving right).
2. Visual Cue: A single RDK was displayed inside the center aperture, which was preceding the stimulus presentation. The horizontal motion direction indicated the target’s horizontal motion direction. The properties of the RDK were matched to those described above for each aperture, but with only horizontal, and no vertical, motion (i.e., no motion transparency).
3. Auditory Cue: A pure 44.1 kHz tone measured at 83 dB at maximum coherence per ear with 50dB background noise, that traveled either from left to right ear (rightward motion) or right to left ear (leftward motion) by fading the volume from one ear to the other (interaural level differences, ILD) was played through headphones in the interval preceding the stimulus presentation. During the tone (the cue) a single blank aperture was presented in the center of the screen for the duration of the cue.
4. Combined Cue: Both the visual cue and the auditory cue as described above were presented simultaneously. Visual cues and auditory cues always had congruent horizontal motion to each other (so that in the invalid cue trials, both cues indicating the incorrect direction of motion).

In order to compare the visual and auditory cues, the salience of the cues was adjusted such that the ability to distinguish the presented horizontal motion direction was consistent for each subject. In order to obtain multiple cue levels (threshold and sub-threshold), we used a two-stage procedure for selecting the appropriate difficulty levels: first, a three-down, one-up adaptive staircase (Vaina et al., 2003) for both visual and auditory cues to determine approximate threshold, then a constant stimulus procedure for precise estimation of the psychometric response function. For the visual cues, difficulty was varied by adjusting the motion coherence (e.g., the proportion of dots moving left/right, with the rest repositioned randomly). For the auditory cues, the extent of the ILD was varied (with 100% coherence represented as a tone that started at 100% volume in the starting ear and 0% volume in the ending ear, and moving to the opposite; for lower coherences, the starting and ending volumes were adjusted such that 0% coherence indicated a sound which did not move at all). In all these conditions, the staircase terminated after 10 reversals, and coherence levels for the last 6 reversals were averaged to obtain a threshold estimate. This adaptive staircase was repeated three times and averaged across all blocks to determine the subject’s threshold detection level. Constant stimulus blocks were then administered with 7 coherence levels (at threshold, along with three levels above threshold and three levels below), with 60 trials per level presented in random order. The accuracy vs coherence curve was fit to a two-parameter sigmoid psychometric function. For each subject, the coherence level that corresponded to 76% accuracy (d’ = 1) was used as a threshold level, and the coherence level that corresponded to 63.8% accuracy (d’ = 0.5) was used as a subthreshold level for each cue.

### Task

Each trial began with a fixation cross (40×40 arcmin; 43.9 cd m^−2^ luminance) displayed in the center of screen. After 300 ms the outline of a single circular aperture was displayed in the center of the screen for 300 ms while the cue was displayed (in the case of no cue, the aperture was displayed for the same 300 ms). This was followed by a 300 ms stimulus onset asynchrony (SOA) showing only the fixation cross in the center of the screen. It was immediately followed by four transparent motion apertures (as described above) displayed for 1000 ms. The subjects performed a dual task, providing two responses on each trial. *Task 1:* subjects were prompted to press a designated key (1-4) on the computer-attached keypad to indicate which of the four quadrants displayed the target aperture (the aperture containing a horizontal RDK with motion opposite the other three). *Task 2:* subjects were asked to report, by pressing the upward or downward arrow on the keypad, the direction of the vertical motion RDK in the target aperture. Performance was evaluated based on the aperture they selected, even if this was not the correct target aperture, that is, if they identified the wrong target aperture, but correctly identified the motion in that aperture, the second response was considered correct since they were able to identify the motion within the aperture being attended. The responses and reaction times were recorded relative to the end of the 1000 ms display of the four apertures containing transparent motion. Subjects had unlimited time to respond to enter both responses (Fig 1), however, they were instructed to respond as quickly as possible and as accurately as possible. There was a 300 ms inter-stimulus interval (ISI) between all trials.

**Figure 1.**
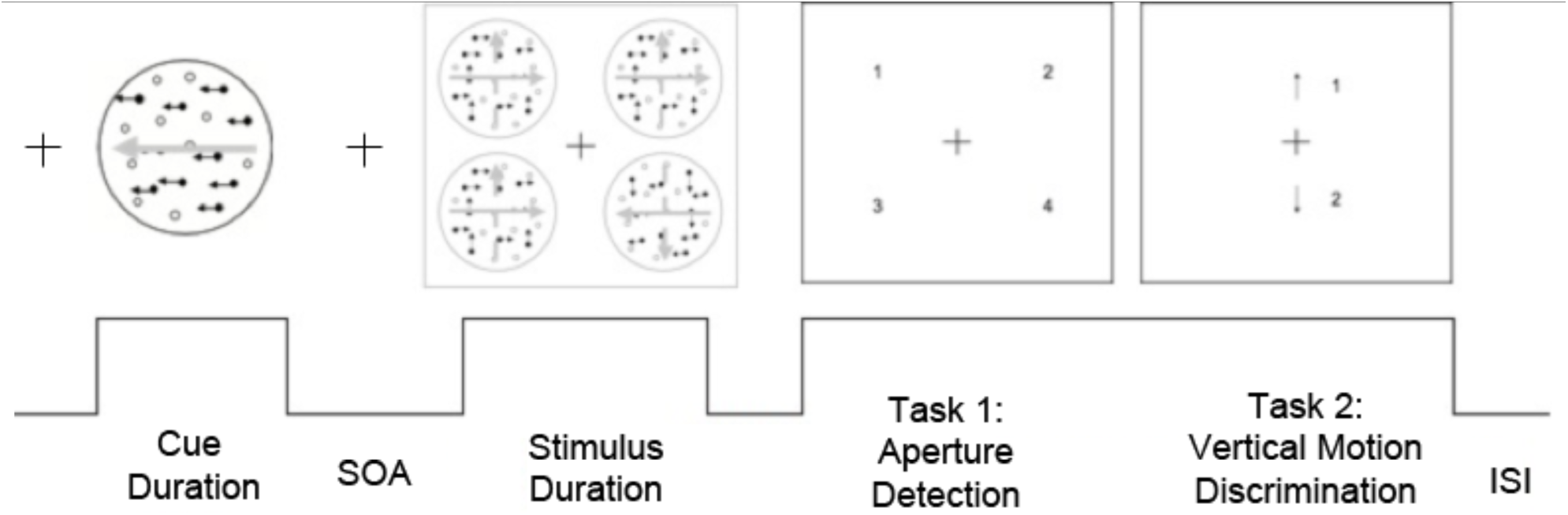
Diagram of a single trial showing the visual cue condition. The cue is presented for 300 ms followed by a 300 ms stimulus onset asynchrony (SOA). Stimulus is then displayed for 1 second. During the response time, in Task 1, observers must detect the target aperture which contains the cued horizontal motion, followed immediately by Task 2, which involves the vertical motion discrimination in the target aperture. There was a 300 ms interstimulus interval (ISI) between all trials.

### Block Types

Data was collected in separate block types for each subject: (1) *Uniform*: 20 trials of each cue type (no cue, visual cue, auditory cue, or combined cue) one at a time. Subjects were told prior to each block which cue type would be provided. (2) *Interleaved:* 20 trials of each of the no cue, visual cue, and auditory cue conditions (60 total trials) were presented in a pseudorandomized interleaved sequence, such that subjects did not know on any given trial which cue type would be presented. The test order of each block type and threshold type was randomized for each subject. All subjects completed the task over two days in separate sessions, with half the total number of blocks in each session.

### Statistical analyses

The way the modality, saliency and consistency of the cues affected subjects’ accuracy in the motion processing tasks were modeled using multifactorial generalized linear mixed effects analyses for binomial dependent variables (GLME; the glmer function of the R lme4 module). Binomial GLMEs were constructed 1) to compared the effect of valid visual, auditory, and audiovisual cues to conditions with no cues, 2) to compare attention performance during tasks with valid vs. invalid attention cues, 3) to examine how the cue salience affects attention performance (Threshold vs. Subthreshold conditions), and 4) to determine the effects of cue predictability (Uniform vs. Interleaved blocks). The results are presented by cue threshold level and by block type. The effects of task parameters on RT performance were analyzed using linear mixed effects models (LME; the lmer function of the R lme4 module). A contrast setting that sums to zero in analyses using the Type III Wald χ^2^ test for hypothesis testing based on our GLME and LME models (Fox, 1997). We used the Anova function in the R car module to determine the statistical significance values in all analyses.

## Results

### Effects of Valid Threshold-Level Visual and Crossmodal Auditory Motion Cues on Motion Processing Accuracy

We first analyzed the effect of intramodal and crossmodal motion cues on the motion aperture detection (Task 1) across all trials within all subjects using a binomial GLME model that examined the fixed effects of the visual cues (on vs. off), auditory cues (on vs. off), and the interaction between visual and auditory cues (nested within the uniform condition only), and which controlled for the fixed effect of the block type (uniform vs. interleaved) as well as the random effect of the subject identity. This analysis showed a highly significant improvement of the target aperture detection by both visual (Wald χ^2^= 18.8, p<0.001) and crossmodal auditory cues (Wald χ^2^ *=* 16.4, p<0.001), in comparison to the no cue condition (Fig. 2A). We then used a similar binomial GLME model to examine the effect of visual and auditory cues on the motion direction discrimination (Task 2; Fig. 2B). A significant improvement of performance accuracy was observed after both visual (Wald χ^2^ *=* 13.3, *p*<0.001) and auditory cues (Wald χ^2^ *=* 7.4, *p*<0.01).

**Figure 2.**
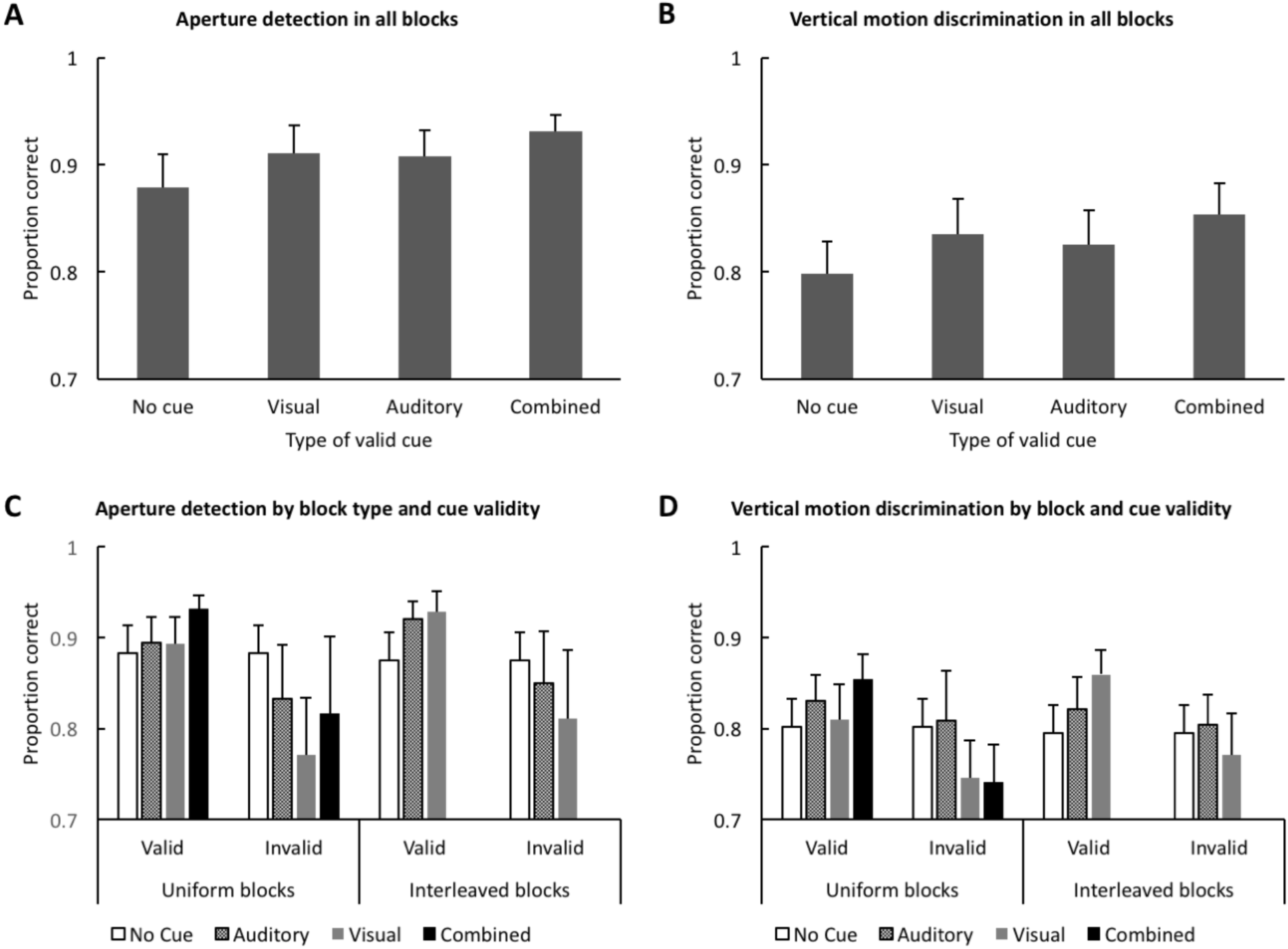
Average motion processing accuracy in the threshold condition across subjects. Error bars reflect the standard error of mean. A) Aperture detection (Task 1) performance accuracy in uniform and interleaved blocks in the threshold cue condition. B) Vertical motion discrimination (Task 2) performance accuracy after valid cues in uniform and interleaved blocks in the threshold cue condition. C) Aperture detection accuracy in different blocks, for valid vs. invalid cues. D) Vertical motion discrimination in different blocks, for valid vs. invalid cues. Taken together, the data show a significant improvement of visual motion processing accuracy after valid intramodal visual, crossmodal auditory, and bimodal visual-auditory cues. On the other hand, the invalid motion cues deteriorated the performance accuracy, which corroborates that the subjects utilized the cues.

Below, we will describe more detailed analyses, which further elucidate the effects of visual and auditory motion cues in the separate task conditions (uniform vs. interleaved block types, threshold vs. subthreshold cues) as well as the effects of cue validity that were examined to verify that the cues, indeed, benefited task performance

RT results, which were generally less significant and clear than the effects observed in accuracy measures, will be described last.

### Effects of Threshold Cues in Uniform Blocks on Motion Processing Accuracy

The effect of threshold cues on motion processing accuracy in uniform blocks are shown in Fig. 2C. To test the statistical significance of these effects, we used a GLME model that predicted the accuracy of aperture detection as a function of the fixed effects of valid visual cues (on or off), valid auditory cues (on or off), and the interaction of valid auditory and visual cues, and the random effect of subject identity. This binomial GLME analysis suggested that the accuracy of aperture detection was significantly improved by valid visual cues (Wald χ^2^ = 7.6, p<0.01) and valid auditory cues (χ^2^= 8.8, p<0.01). There was only a non-significant trend toward an interaction between the visual and auditory cues (χ^2^=3.0, p=0.08), suggesting that the effect of multisensory attention cueing is, for the most part, explainable by the additive effects of unimodal visual and auditory cues. To verify that adding auditory information improved performance, we also computed a specific contrast between the unimodal and bimodal cue conditions. This analysis showed that the aperture detection accuracy was significantly better in the bimodal condition than in the unimodal visual cue condition (Wald χ^2^ = 9.6, p<0.01) and in the unimodal auditory cue condition (Wald χ^2^= 8.5, p<0.01).

Robust support for the interpretation that the subjects attended to and relied on the cues, was obtained from GLMEs comparing the effects of valid and invalid cue conditions (Fig. 2C). The effects of cue validity were tested using binomial GLME analyses, which modeled the accuracy of aperture detection as a function of cue type (visual, auditory, or audiovisual), cue validity (valid or invalid), and the interaction of cue type and validity, and which controlled for the random effects of subject identity. The results suggested a highly significant effect of cue validity, irrespective of the cue modality (Wald χ^2^= 61.0, p<0.001).

The results of more detailed analysis of vertical motion direction accuracy (Task 2) for the threshold cues are shown in Fig. 2D. We analyzed how the valid cues affected this task performance in different task blocks using a binomial GLME analysis, which modeled the accuracy of vertical motion detection as a function of the fixed effects of valid visual cues, valid auditory cues, and the interaction across visual and auditory cues, and which controlled for the random effects of subject identity. The results of this GLME model suggested a significant main effect of auditory cue (Wald χ^2^= 9.4, p<0.01).

To verify that the subjects benefited from the cues, we also used a binomial GLME analysis, which modeled the accuracy of vertical motion direction discrimination as a function of the fixed effects of the cue type (visual, auditory, or audiovisual), the cue validity (valid or invalid), and the interaction of the cue type and validity, and which controlled for the random effects of subject identity. This GLME model showed a highly significant effect of cue validity (Wald χ^2^= 17.6, p<0.001), consistent with the descriptive results shown in Fig. 2D.

### Comparison of Motion Processing Accuracy in Uniform vs. Interleaved Blocks

In the uniform blocks, the cue modality was fixed throughout each testing block so that subjects knew which cue will be presented on each trial and, thus, they could allocate attention to that modality alone. To control for the effects of splitting attention among the visual and auditory modalities, and the impact of switching between cue modalities, we compared the effects of visual and auditory cues, as well as the effect cue validity, across the uniform and interleaved task conditions for the threshold-level cues (Fig. 2C). Overall, it seemed that the cue switching across cue modalities did not significantly decrease the aperture detection accuracy. To verify this statistically, we used a binomial GMLE, which modeled the trial-to-trial accuracy of aperture detection as a function of the fixed effects of the cue type (visual or auditory), cue validity (valid or invalid), the block type (uniform vs. interleaved), and the interactions across these fixed factors and which controlled for the random effects of subject identity. This GLME showed a highly significant main effect of cue validity (Wald χ^2^= 69.4, p<0.001) but no evidence for an interaction between the cue validity and block type or between the cue type and block type. We also verified the effect of valid visual and valid auditory cues in the interleaved condition using more specific binomial GLME contrasts. Valid cues led to a highly significant improvement in performance for both visual (Wald χ^2=^ 21.5, p<0.001) and auditory (Wald χ^2^ = 16.2, p<0.001) cues compared to the no cue condition. Taken together, these results suggest that the effect of auditory and visual cues as well as valid vs. invalid cues was consistent, independent of the predictability of cue type (see Fig. 2C).

We used analogous binomial GLME models to examine accuracy of in the direction discrimination task (Fig. 2D). A binomial GMLE, which modeled the trial-to-trial accuracy of motion direction discrimination as a function of the fixed effects of the cue type (visual or auditory), cue validity (valid or invalid), the block type, and the interactions between these fixed factors, and which controlled for the random effects of subject identity. This GLME showed a significant main effect of cue validity (Wald χ^2^ = 12.0, p<0.01) but no evidence for an interaction between the cue validity vs. block type/event predictability or the cue type vs. block type. The more specific more specific binomial GLME contrasts showed a significant improvement of performance by visual cues (Wald χ^2^= 20.2, p<0.001), but the effect of auditory cues did not quite reach statistical significance (Wald χ^2^= 3.1, p=0.08) cues compared to the no cue condition.

### Effect of Cue Salience on Motion Processing Accuracy in Uniform Blocks

To analyze the effect of cue salience, we used a binomial GLME that modeled the accuracy of aperture detection as a function of the fixed effects of cue salience (threshold, i.e., d’=1 vs. subthreshold, i.e., d’=0.5), cue type (visual, auditory, audiovisual), cue validity (valid vs. invalid), and the interaction between the cue type and validity, while controlling for the random effect of subject identity. As expected, the effect of cueing was significantly more beneficial in the threshold vs. subthreshold conditions, shown by a main effect of cue salience (Wald χ^2^= 4.7, p<0.05). As a matter of fact, in the subthreshold condition, particularly the hard-to-detect auditory cues seemed to distract the aperture detection. However, even in the case of subthreshold cues, the accuracy of aperture detection was significantly better when the cues were valid vs. invalid (Wald χ^2^= 12.3, p<0.001). The effect of cue salience was analogous in the vertical motion direction task, which were analyzed using binomial GLMEs analogous to those used for the aperture detection task. There were significant main effects of cue salience (Wald χ^2^= 6.9, p<0.01) and validity (Wald χ^2^ = 19.2, p<0.001) in the main GLME that considered the fixed effects of salience, cue type, validity, and the interaction between cue type and validity, as well as the random effect of subject identity. Further, the effect of cue validity was highly significant also within the blocks with subthreshold cues only (Wald χ^2^= 4.2, p<0.05).

### Reaction Times (RT) in Threshold Cue Conditions

RT data for the aperture detection task is described in **Table 1**. In the uniform blocks, no significant effects of valid visual, auditory, or audiovisual threshold cues were observed in our LME models. However, consistent with the accuracy effects, a LME analysis that modeled RT in the aperture detection task as a function of cue validity and type, as well as their interaction, and controlled for the random effect of subject, revealed that RTs were significantly faster for valid than invalid cues (Wald χ^2^=10.6, p<0.01).

**Table 1.** Aperture RT data in different task conditions. For this table, the trial-specific RT values were first aggregated within subjects, after which the group mean and standard deviations (SD) were calculated.

| **Salience** | **Block type** | **Validity** | **Cue type** | **Mean** | **SD** |
| --- | --- | --- | --- | --- | --- |
| Threshold | Uniform | Valid | A | 0.13 | 0.26 |
|  |  |  | V | 0.15 | 0.30 |
|  |  |  | VA | 0.13 | 0.33 |
|  |  | Invalid | A | 0.21 | 0.36 |
|  |  |  | V | 0.27 | 0.40 |
|  |  |  | VA | 0.28 | 0.46 |
|  | Interleaved | Valid | A | 0.16 | 0.30 |
|  |  |  | V | 0.10 | 0.32 |
|  |  | Invalid | A | 0.20 | 0.34 |
|  |  |  | V | 0.24 | 0.33 |
| Subthreshold | Uniform | Valid | A | 0.22 | 0.38 |
|  |  |  | V | 0.12 | 0.24 |
|  |  |  | VA | 0.11 | 0.27 |
|  |  | Invalid | A | 0.25 | 0.36 |
|  |  |  | V | 0.11 | 0.30 |
|  |  |  | VA | 0.19 | 0.37 |
|  | Interleaved | Valid | A | 0.18 | 0.27 |
|  |  |  | V | 0.11 | 0.27 |
|  |  | Invalid | A | 0.13 | 0.23 |
|  |  |  | V | 0.13 | 0.23 |
| No cue | Uniform |  |  | 0.14 | 0.29 |
|  | Interleaved |  |  | 0.19 | 0.31 |

In the interleaved condition, there was a significant main effect of cue type (Wald χ^2^=14.4, p<0.001) in a LME analysis that modeled RT in the aperture detection task as a function of valid cues (visual vs. auditory vs. no cue) and the random effect of subject identity. According to more detailed comparisons between each cue type and no cue conditions, visual cues decreased the RT in the aperture detection task significantly (χ^2^=13.8, p<0.001), but no significant effects for the auditory cue were observed. Similar to the uniform blocks, RTs were significantly faster for valid vs. invalid cues (χ^2^=7.4, p<0.01).

RT data in the direction discrimination task is described in Table 2. A weakly significant increase of RTs was observed after valid visual cues in the uniform condition (χ^2^=4.0, p<0.05). No other statistically significant effects of cueing or cue validity were observed in analyses on RTs in this task.

**Table 2.** Direction RT data in different task conditions. For this table, the trial-specific RT values were first aggregated within subjects, after which the group mean and standard deviations (SD) were calculated.

| Saliency | Block type | Validity | Cue type | Mean | SD |
| --- | --- | --- | --- | --- | --- |
| Threshold | Uniform | Valid | A | 0.35 | 0.13 |
|  |  |  | V | 0.36 | 0.14 |
|  |  |  | VA | 0.37 | 0.17 |
|  |  | Invalid | A | 0.36 | 0.16 |
|  |  |  | V | 0.35 | 0.16 |
|  |  |  | VA | 0.35 | 0.16 |
|  | Interleaved | Valid | A | 0.36 | 0.13 |
|  |  |  | V | 0.36 | 0.14 |
|  |  | Invalid | A | 0.36 | 0.16 |
|  |  |  | V | 0.38 | 0.14 |
| Subthreshold | Uniform | Valid | A | 0.37 | 0.16 |
|  |  |  | V | 0.35 | 0.15 |
|  |  |  | VA | 0.38 | 0.16 |
|  |  | Invalid | A | 0.33 | 0.10 |
|  |  |  | V | 0.35 | 0.15 |
|  |  |  | VA | 0.36 | 0.19 |
|  | Interleaved | Valid | A | 0.36 | 0.15 |
|  |  |  | V | 0.35 | 0.12 |
|  |  | Invalid | A | 0.36 | 0.13 |
|  |  |  | V | 0.35 | 0.15 |
| No cue | Uniform |  |  | 0.34 | 0.14 |
|  | Interleaved |  |  | 0.35 | 0.14 |

### RTs in Subthreshold Cue Conditions

In the uniform blocks, a LME model that considered the fixed effects of visual and auditory cues, as well as their interactions, and controlled for the random effects of subject identity suggested that valid visual cues significantly improved RTs performance in the aperture detection task (Wald χ^2^=11.4, p<0.001). More specific contrasts, further, suggested that in this task condition, valid auditory cues had a distracting effect that increased RTs of the aperture detection (Wald χ^2^=6.17, p<0.05). The effects of cue validity were non-significant.

In the interleaved condition, there was a significant main effect of cue type (Wald χ^2^=11.6, p<0.01) in a LME analysis that modeled RT in the aperture detection task as a function of valid cues (visual vs. auditory vs. no cue) and the random effect of subject identity. Specific comparisons to the no cue conditions suggested that visual cues decreased RTs in the aperture detection task significantly (Wald χ^2^=11.4, p<0.001), but no significant effects for the auditory cue were observed. No significant RT effects of cue validity were observed.

No statistically significant RT improvements were observed in the direction discrimination task in our LME models in any of the subthreshold conditions. In contrast, similarly to the aperture detection task, it seemed like the auditory cues resulted in a distracting effect, reflected by a RT delay (Wald χ^2^=5.8, p<0.05).

## Discussion

In this study, observers were presented with visual, auditory, or combined visual-auditory feature cues implementing horizontal direction information and were asked to identify a target aperture from among four transparent motion apertures, one of which contained a horizontal motion component moving in the direction of the cue. Importantly, the target aperture could be identified in the absence of any cue as the oddball motion target (i.e., the single leftward moving target among 3 other rightward moving targets). This suggests that the cue was not necessary to performing the task, yet it could provide a prior for motion direction that would facilitate detection of the target aperture. In addition to the horizontal motion, all four apertures contained an overlapping vertical motion dot field and subjects were asked to detect the direction of vertical motion in the aperture they chose to be the target.

By matching the salience (visual motion coherence, auditory amplitude fade) of visual and auditory cues, we measured how the specific cue modality affected a subject’s ability to use feature-based attention to the direction of motion to facilitate target detection. Using this approach, we found that valid visual, auditory, and audiovisual cues reliably showed an improvement in target detection when presented at threshold levels. On the other hand, our analyses suggested that cues were overall significantly more beneficial in the threshold than subthreshold conditions. These results extend previous knowledge of how cross-modal cueing affects target detection: Previous studies have examined to which degree cross-modal cues affect spatial attention in different modalities (Spence et al., 1998, Driver and Spence, 1998), but they have not adjusted the cue features themselves to measure the extent of usefulness. Others (Pessoa et al., 2009) have adjusted the color features of cues, and used the cues for matching a grating orientation, but have not investigate the ability of such cross modal cueing to facilitate motion processing.

### Valid vs. Invalid Cues

One of the most prevalent effects observed throughout the task was a significant improvement of performance accuracy in the valid cue trials vs. invalid cue trials. At the threshold cue coherence, the invalid cues of all types caused worse performance on aperture discrimination than valid cues. This valid vs. invalid cue performance difference provides a reliable indication of whether or not subjects are attending to and using the cue when attempting to determine the target aperture. At the same time, this performance decrease mirrors the results of (Posner and Peterson, 1980) in which reaction times were slower following invalid cues than valid cues. Similarly, our results showed evidence for both increased reaction times (aperture detection) and decreased performance with invalid cue types. This supports the attentional shift paradigm in which resources are allocated for efficient processing when valid cues are used. With the invalid cues, there may be a cost to performance as there is an increased effort added to shifting attention. Because of the difficulty in this motion perception task and the limited time of stimulus presentation, the extra attentional shift diverted to invalid cues might have been enough to significantly reduce performance.

### Cross-modal Attention

In our study, when looking at cross-modal effects, we also found that apart from the subthreshold condition, in which the cue processing could have had a distractor effect, auditory cues alone were generally as effective as visual cues. The beneficial effects of auditory cues were greatest during interleaved runs, which were overall more difficult than the uniform blocks. However, in the uniform blocks, the combination of both visual and auditory cues tended to have the strongest positive effects on performance for both the target aperture detection task and for the vertical motion direction discrimination task.

Additive multisensory effects have been shown in previous studies with focus on audiovisual integration (Burr and Alais, 2006). It has been, for example, reported that visual and auditory coherence combined have a summation effect with statistical optimal combination of signals based on maximum likelihood estimation with an increase in performance with a factor of about 1/√2. Our results, which suggested only non-significant trends toward visual-auditory interactions in attentional cueing, are seemingly consistent with these studies. However, it has to be noted that the present study did not measure detection of the motion cues themselves.

In conclusion, whereas previous studies have shown crossmodal effects of spatial attention, our results demonstrate a spread of crossmodal feature-based attention cues, which have been matched for the detection threshold, on visual target detection. These effects were evident in comparisons between cued and uncued conditions, as well as in tasks that compared the effects of valid vs. invalid cues.

## Acknowledgements

This work was supported by the National Science Foundation grant 1545668 (LMV), and by the National Institutes of Health grant R21DC014134 (JA).

